# gwasrapidd: an R package to query, download and wrangle GWAS Catalog data

**DOI:** 10.1101/643940

**Authors:** Ramiro Magno, Ana -Teresa Maia

**Affiliations:** Centre for Biomedical Research (CBMR), Universidade do Algarve, Gambelas Campus, 8005-139 Faro, Portugal; Algarve Biomedical Center (ABC), Universidade do Algarve, Gambelas Campus, 8005-139 Faro, Portugal; Departamento de Ciências Biomédicas e Medicina (DCBM), Universidade do Algarve, Gambelas Campus, 8005-139 Faro, Portugal

**Keywords:** Genome-wide association studies, GWAS, disease susceptibility, Genomics, GWAS Catalog, R, SNP, trait, trait-ontology, Software, human, REST client

## Abstract

**Motivation:** The NHGRI Catalog of Published Genome-Wide Association Studies (GWAS) Catalog has collected, curated, and made available data from over 3 900 studies. The recently developed GWAS Catalog REST API is the only method allowing programmatic access to this resource.

**Results:** Here, we describe *gwasrapidd*, an R package that provides a client interface to the GWAS Catalog REST API, representing an important software counterpart to the server-side component. *gwasrapidd* enables users to quickly retrieve, filter and integrate data with comprehensive bioinformatics analysis tools, which is particularly critical for those looking into functional characterisation of risk loci.

**Availability:** *gwasrapidd* is freely available under an MIT License, and can be accessed from https://github.com/ramiromagno/gwasrapidd.

## Introduction

Over the last 15 years, genome-wide association studies (GWAS) have greatly extended our knowledge of the genetic risk for human complex diseases, and have provided starting points for the understanding of the underlying disease mechanisms, raising possibilities for use in clinical patient/general population stratification [1].

The National Human Genome Research Institute (NHGRI) Catalog of Published GWAS Catalog, created in 2014, is a publicly available, manually curated, database of all published GWA studies [2]. Its latest data release [date 2019-05-03] includes data from 3,989 publications and 138,312 unique SNP-trait associations for human diseases. Currently, these data can be accessed by three methods: (i) via the web graphical user interface, (ii) by downloading database dumps, or, more recently, (iii) via the GWAS Catalog representational state transfer (REST) application programming interface (API).

Hitherto, the only software that provides parsing of GWAS Catalog data into R is the Bioconductor package *gwascat* [3], which uses the aforementioned database dumps to read in the data. However, the REST API is the only method allowing direct programmatic access, and hence the preferred method for bioinformatics analyses. Yet, so far, no software solution exists to ease the querying and parsing of data returned by it.

To address this limitation, we have developed an R package [4] that provides programmatic access to the GWAS Catalog REST API: *gwasrapidd*. This package provides a simple interface for querying Catalog data, abstracting away the informatic details of the REST API. In addition, retrieved data is mapped to in-memory relational databases of tidy data tables, allowing prompt integration with tidyverse packages for subsequent transformation, visualisation and modelling of data [5, 6].

## Results

### Retrieving data from the GWAS Catalog REST API

The GWAS Catalog REST API is an EBI service hosted at https://www.ebi.ac.uk/gwas/rest/api/. The REST API uses hypermedia with resource responses following the JSON Hypertext Application Language (HAL) format [7]. Response data is therefore provided as hierarchical data in JSON format. Although flexible, hierarchical data is not straightforwardly convertible to tabular format — the preferred format for data analysis in R [6]. Moreover, a resource response can be paginated, i.e., split into multiple responses, requiring navigation over all the individual responses, requiring posterior aggregation. Finally, data can also be embedded, i.e., have other resources contained within them, adding extra complexity to the returned JSON format (Additional file 1: Table S1, and [8]).

To ease the conversion from the hierarchical to the relational tabular format, and to abstract away the informatic details associated with the HAL format, we developed a set of retrieval functions (Fig. 1A). Since the REST API data is organised around four core data entities —*studies, associations, variants* and *traits* [8]— we implemented four corresponding retrieval functions that encapsulate the technical aspects of resource querying and format conversion: `get studies, get associations, get variants and get traits` (Fig. 1A). These functions simplify the querying of GWAS entities, by providing a complete and consistent interface to the Catalog. For example, to query for *studies*, the user needs only to know the function `get studies`, whereas the REST API itself exposes a set of disparate resource URL endpoints for *studies* following the available search criteria (Additional file 1: Table S1, and [8]). Moreover, the user can choose from any number of available search criteria exposed by the REST directly as arguments to the retrieval functions (Fig. 1B). All arguments are vectorised, meaning that multiple queries are promptly available from a single function call. Results obtained from multiple queries can be combined in an `OR` or `AND` fashion with the `set operation` parameter. If `set operation` is set to `OR` (default behaviour), results are collated while removing duplicates, if any. If `set operation` is set to `AND`, only entities that concomitantly match all criteria are returned. If finer control is needed on combining results from different queries, the user can make separate calls with the retrieval functions and combine the results from the different objects with any combination of these functions: `bind(), union(), intersect(), setdiff()` and `setequal()`.These are S4 methods that work with the S4 classes created in *gwasrapidd* (Fig. 2).

**Figure 1.**
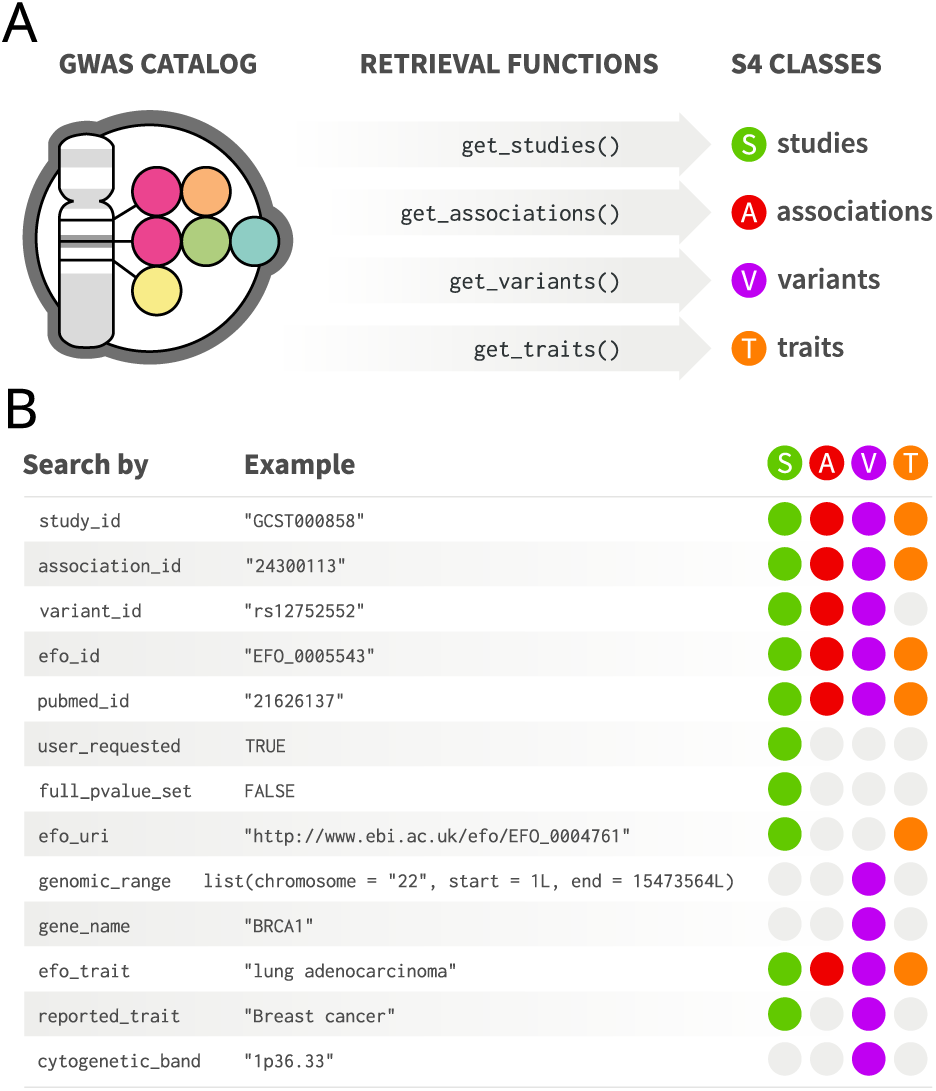
*gwasrapidd* retrieval functions. **(A)** Functions for retrieving data from the GWAS Catalog: `get studies(), get associations(), get variants()` and `get traits()`. **(B)** *gwasrapidd* search criteria (function parameters) to be used with retrieval functions. Coloured circles indicate which entities can be retrieved by which criteria.

**Figure 2.**
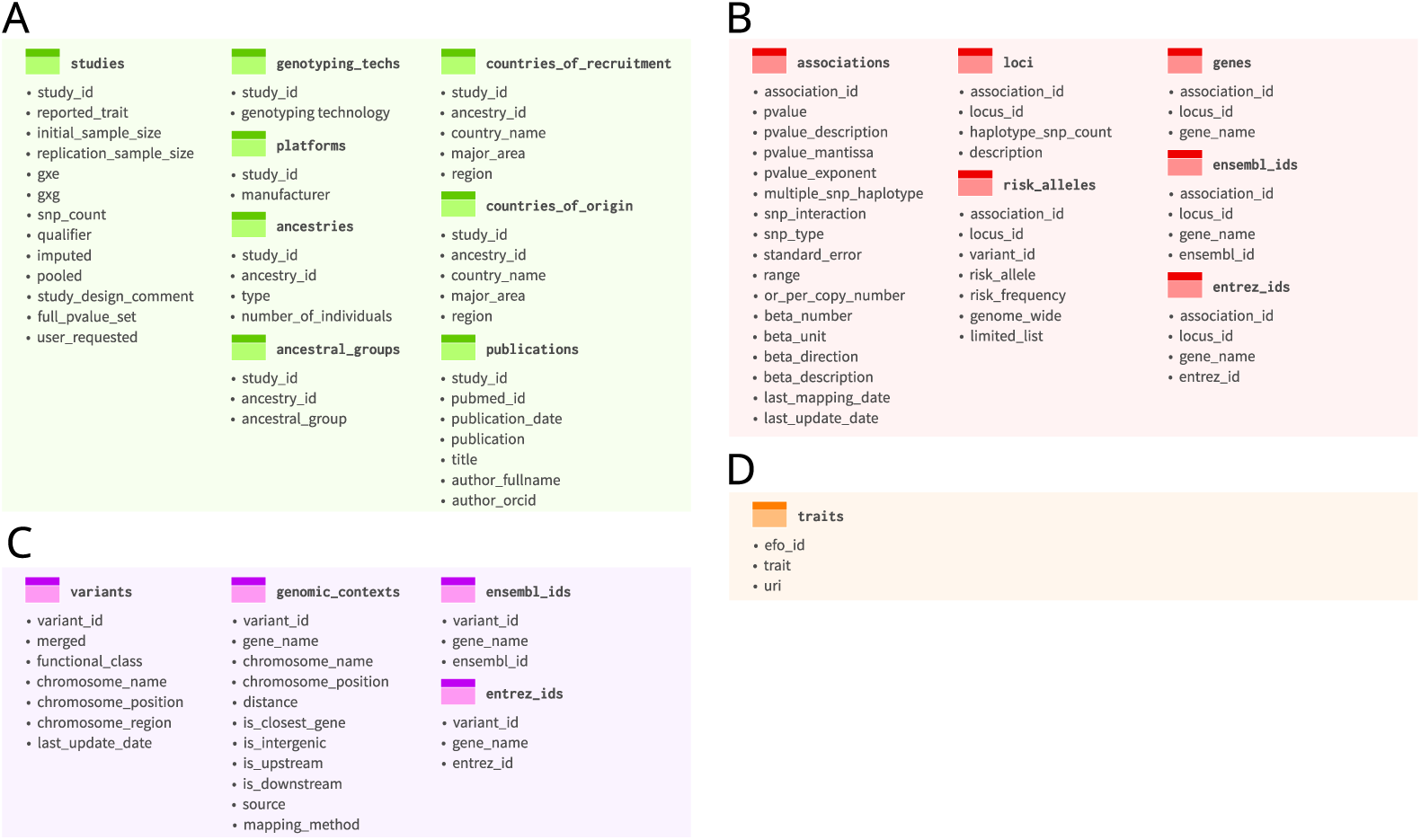
S4 representation of *studies, associations, variants* and *traits*. **(A)** S4 studies object, comprising 8 slots, i.e., 8 tables: `studies, genotyping techs, platforms, ancestries, ancestral groups, countries of recruitment, countries of origin` and `publications`. All tables share the primary key `study id`. **(B)** S4 `associations` object, comprising 6 slots, i.e., 6 tables: `associations, loci, risk alleles, genes, ensembl ids` and `entrez ids`. All tables share the primary key `association id`. **(C)** S4 `variants` object, comprising 4 slots, i.e., 4 tables: `variants, genomic contexts, ensembl ids` and `entrez ids`. All tables share the primary key `variant id`. **(D)** S4 `traits` object, with one slot, i.e., one single tables: `traits`. The `traits` has the primary key `efo id`.

### Representation of GWAS Catalog entities

The GWAS Catalog REST API provides the data organised according to four key concepts: *studies, associations, variants* and *traits*; reflected on the possible types of JSON responses. In *gwasrapidd*, each of these four types of responses to S4 objects are mapped to four classes of the same name: `studies, associations, variants` and `traits`, respectively (Fig. 2). All S4 classes share the same design principles that makes them relational databases: (i) each slot corresponds to a table (dataframe in R), (ii) the first slot corresponds to the main table that lists observations of the respective GWAS Catalog entity, e.g., *studies*, and, (iii) all tables have a primary key, the identifier of the respective GWAS Catalog entity: `study id, association id, variant id` or `efo id` (Fig. 2). For easy consultation of the variables in the tables, we provide a cheatsheet (Additional file 2: gwasrapidd cheatsheet); for the detailed description the user can issue the following commands to open the help page about each class: `class?studies, class?associations, class?variants or class?traits`.

### Case study

To illustrate how to use *gwasrapidd*, we take as a motivating example the work by Light *et al.* [9]. In this work the authors aimed at characterising chromatin status at allelic resolution, as a strategy for elucidanting the cis-regulatory mechanisms behind complex disease risk. To perform this characterisation, the authors started by selecting variants previously reported in the GWAS Catalog for autoimmune disease. We enact now what could have been these first steps of their approach using *gwasrapidd*. We start by loading *gwasrapidd* :

**library** (gwasrapidd)

Then following by querying the GWAS Catalog for *studies* by searching by *autoimmune disease* (an EFO trait):

~~~
my_studies <-
get_studies(efo_trait = ‘autoimmune disease’)
~~~

One can now check how many GWAS studies were retrieved using function n(). The same function could be used for the other entities: `associations, variants or traits`.

~~~
n(my_studies)
#> [1] 1
my_studies@studies$study_id
#> [1] “GCST003097”
~~~

Seemingly, only one study matched exactly `‘autoimmune disease’` : the study with the identifier `GCST003097`. We can inspect the original publication(s) that underlie this GWA study entry in the Catalog. For example, to access the associated publication title one can access the `title` variable from the `publications` table:

~~~
my_studies@publications$title
#> [1] “Meta-analysis of shared genetic architecture across ten pediatric
*c→* autoimmune diseases.”
~~~

To quickly browse to the PubMed entry for this publication, the user may use the helper function `open in pubmed()`:

~~~
# This launches your web browser at https://www.ncbi.nlm.nih.gov/pubmed/26301688
open_in_pubmed(my_studies@publications$pubmed_id)
~~~

Now if we want to know the variants previously associated with autoimmune disease, as used by Light et al. (2014), we need to retrieve statistical association information on these variants, and then apply a filter using the same level of significance *P* < 1 × 10^−6^ [9]. A quick inspection at the *gwasrapidd* cheatsheet (Additional file 2: gwasrapidd cheatsheet) hints that statitiscal information, such as p-values and odds ratios can be found in the associations table of the associations S4 class object. So, we can get the associations by the previously found *study* identifier (GCST003097):

~~~
# Alternative query that would work too:
# get_associations(
# efo_trait = ‘autoimmune disease’
# )
my_associations <-get_associations(study_id = “GCST003097”)
~~~

We find 46 *associations* :

~~~
n(my_associations)
#> [1] 46
~~~

However, it might be that not all variants meet the level of significance, as required by Light et al. (2014). We use now functions from the *dplyr* package [10] to filter the table `associations` from the S4 object `associations` based on the p-value (pvalue column).

~~~
# Get association ids
# for which pvalue is less than 1e-6.
dplyr::filter(
my_associations@associations,
pvalue < 1e-6)%>%
dplyr::pull(association_id) ->
association_ids
~~~

Having the *association* identifiers (`association ids`) that meet the requirement P < 1 ×10^−6^, we can easily create a new S4 object (`my associations2`) containing only those *associations* using the subset operator ‘[’:

~~~
# Extract associations by association id
my_associations2 <-my_associations[association_ids]
~~~

The subset operator ‘[’ can also be used with `integer` values to subset by position. Note that both ways of subsetting will filter all tables within the S4 object for the matched identifiers. Now we can check how many associations are still present in my `associations2`:

~~~
n(my_associations2)
#> [1] 44
~~~

Of the 46 associations found in GWAS Catalog, 44 meet the p-value threshold of 1× 10^−6^. Finally, to see variants, we just need to access the table `risk alleles` from the  `my associations2` object. Here are the variant identifers, risk alleles and risk frequency:

~~~
my_associations2@risk_alleles[**c**(‘variant_id’, ‘risk_allele’, ‘risk_frequency’)]
#> # A tibble: 44 × 3
#>      variant_id risk_allele risk_frequency
#>      <chr>      <chr>     <dbl>
#> 1  rs2066363m    C        0.34
#> 2  rs114846446   A        0.01
#> 3  rs7672495     C        0.18
~~~

## Discussion

We have developed an R client to the GWAS Catalog REST API, thus allowing programmatic access to the database. The main features of *gwasrapidd* are: (i) abstracting away the REST API informatic details by providing a simple and consistent interface, and (ii) a tidy data representation of the GWAS entities, i.e., of *studies, associations, variants* and *traits* in the form of in-memory relational databases. Below, we highlight some of the improvements and limitations of our package.

### Improvements

Compared to the exposed REST API, we have augmented the search possibilities in *gwasrapidd* in two ways: (i) by allowing the user to search for *variants* by cytogenetic region (as is possible with the web GUI), and (ii) by allowing searching *variants* by EFO identifier (efo id). Searching by cytogenetic region was implemented by embedding a dataset of genomic ranges of the human cytogenetic bands in *gwasrapidd*. In this way, queries made with cytogenetic band names can be translated to genomic ranges, which are then used with the exposed interface of the REST API to search by genomic range (`genomic range`). Searching *variants* by EFO identifier (`efo id`) is not directly available in the current version of the REST API. Therefore, to create this possibility in *gwasrapidd*, we map EFO identifiers (`efo id`) to EFO traits with `get traits ()` using the former as a search parameter. The table `traits` of the S4 object `traits` contains the `trait` column, providing a one-to-one mapping between each EFO identifier and its trait description (`trait`). Then, we use internally `get variants ()` to search by EFO trait description (`efo trait`).

Finally, *gwasrapidd* also provides a set of helper functions to easily browse linked web resources, such as PubMed [11], dbSNP [12],and GTEx project [13].

## Limitations

The most popular method of access to the Catalog is still the web graphical user interface (GUI), and is therefore still the one that gets more attention and dedication from the GWAS Catalog team. Thus, compared to the web GUI, the REST API is still lagging behind in functionality. Here we discuss some of the limitations of the REST API compared to the web GUI.

When searching by publication related information, with the REST API, we can only use the PubMed identifier as a query. Thus, to search using other publication related criteria, such as words in the title or date of publication, one needs to download all studies first with `get studies` and then perform filtering based on variables from the `publications` table of the `studies` object; thus, albeit possible, this introduces unnecessary delays impacting the user experience. Using the web GUI one can promptly use the free text search bar to query publication titles, or dates.

Another important type of query that benefits from the free text search bar is searching by trait. Currently, with the REST API one can search by EFO identifier (`efo id`) or by EFO trait description (`efo trait`) with all retrieval functions (Fig. 1B). The author’s reported trait (`reported trait`) can only be used to retrieve *studies* or *variants*. In both cases, the search terms need to be exact matches. Conversely, with the web GUI one can use free text to get first a list of possible traits and then find other entities such as *studies* and *associations* using the EFO trait of interest. Again, this is still possible with *gwasrapidd*, but indirect. One can retrieve all traits first with `get traits` and then filter by the `trait` using regular expressions to get all traits related to a specific term or collection of terms, and then use any of the four retrieval functions with the `efo trait` parameter. The web GUI also allows to include child trait terms when searching by trait. This is not currently possible with the REST API.

Another limitation with the REST API is that its database release date is older than the web GUI version. As of this writing, the REST API provided data was last updated in 2019-01-12, whereas the web GUI data is dated 2019-04-21. The REST API database release version can be retrieved with the function `get metadata()`.

## Conclusions

We have developed the first R client to the GWAS Catalog REST API, thus greatly facilitating programmatic access to the database. This improves data mining from within R, accelerating the integration of GWAS data into further genomic and biomedical/clinical studies. *gwasrapidd* is freely available under the MIT Licence and can be accessed from https://github.com/ramiromagno/gwasrapidd.

## Availability and requirements

- **Project name**: gwasrapidd.
- **Project home page**: https://rmagno.eu/gwasrapidd.
- **Operating system(s)**: Platform independent.
- **Programming language**: R.
- **Other requirements**: None.
- **License**: MIT.
- **Any restrictions to use by non-academics**: None.

## Funding

This work was supported by national Portuguese funding through FCT—Fundaçaõ para a Cîencia e a Tecnologia and CRESC ALGARVE 2020: UID/BIM/04773/2013 “CBMR”, POCI-01-0145-FEDER-022184 “GenomePT” and ALG-01-0145-FEDER-31477

“DEvoCancer”.

## Author’s contributions

RM devised and wrote the package, and wrote the manuscript. ATM supervised the project and wrote the manuscript. Both authors read and approved the final manuscript.

## Acknowledgements

We would like to thank the GWAS Catalog team, particularly Daniel Suveges, for all the help and support with the REST API throughout the entire development of *gwasrapidd*. We thank also the remaining of the Cancer Functional Genomics lab members for feedback on the user experience of *gwasrapidd*.

## Additional Files

### Additional file 1 — Table S1. GWAS Catalog REST API URL endpoints

- **File name**: `additional file 1.pdf`.
- **File format**: Portable Document Format (PDF).
- **Title**: Table S1. GWAS Catalog REST API URL endpoints.
- **Description**: Additional file 1 contains Supplementary Table 1. Table S1 describes the GWAS Catalog REST API endpoints, along with its parameters (search criteria), and attainable GWAS entities: *studies, associations, variants* and *traits*.

### Additional file 2 — gwasrapidd cheatsheet

- **File name**: `additional file 2.pdf`.
- **File format**: Portable Document Format (PDF).
- **Title**: gwasrapidd cheatsheet.
- **Description**: Additional file 2 contains an infographics: gwasrapidd cheatsheet.

