## Additional File 1 for "gwasrapidd: an R package to query, download and wrangle GWAS Catalog data"

| Search criterion | Examples | Studies | Associations | Variants | Traits |
|---|---|---|---|---|---|
| Study identifier | E□ P□ Studies: /studies/GCST001085/<br>E■ P□ Associations: /studies/GCST001085/associations<br>E■ P□ Variants: /studies/GCST001085/snps<br>E■ P□ Traits: /studies/GCST001085/efoTraits | ■ | ■ | ■ | ■ |
| Association identifier | E□ P□ Studies: /associations/24299710/study<br>E□ P□ Associations: /associations/24299710/<br>E■ P□ Variants: /associations/24299710/snps<br>E■ P□ Traits: /associations/24299710/efoTraits | ■ | ■ | ■ | ■ |
| Variant identifier | E■ P□ Studies: /singleNucleotidePolymorphisms/rs3798440/studies<br>E■ P□ Associations: /singleNucleotidePolymorphisms/rs3798440/associations<br>E□ P□ Variants: /singleNucleotidePolymorphisms/rs3798440/<br>Traits: NA | ■ | ■ | ■ | □ |
| EFO trait identifier | E■ P□ Studies: /efoTraits/EFO_0000537/studies<br>E■ P□ Associations: /efoTraits/EFO_0000537/associations<br>Variants: NA<br>E□ P□ Traits: /efoTraits/EFO_0000537/ | ■ | ■ | □ | ■ |
| PubMed identifier | E■ P■ Studies: /studies/search/findByPublicationIdPubmedId?pubmedId=21626137<br>E■ P□ Associations: /associations/search/findByPubmedId?pubmedId=21626137<br>E■ P■ Variants: /singleNucleotidePolymorphisms/search/findByPubmedId?pubmedId=21626137<br>E■ P■ Traits: /efoTraits/search/findByPubmedId?pubmedId=21626137 | ■ | ■ | ■ | ■ |
| Whether studies were requested by users of the GWAS Catalog or not | E■ P■ Studies: /studies/search/findByUserRequested?userRequested=false<br>Associations: NA<br>Variants: NA<br>Traits: NA | ■ | □ | □ | □ |
| Whether studies have full summary statistics or not | E■ P■ Studies: /studies/search/findByFullPvalueSet?fullPvalueSet=true<br>Associations: NA<br>Variants: NA<br>Traits: NA | ■ | □ | □ | □ |
| EFO URI | E■ P■ Studies: /studies/search/findByEfoUri?uri=http://www.ebi.ac.uk/efo/EFO_0004761<br>Associations: NA<br>Variants: NA<br>E■ P■ Traits: /efoTraits/search/findByEfoUri?uri=http://www.ebi.ac.uk/efo/EFO_0004761 | ■ | □ | □ | ■ |
| Variant and study identifiers concomitantly | Studies: NA<br>E■ P■ Associations: /associations/search/findByRsIdAndAccessionId?rsId=rs16956936&accessionId=GCST000392<br>Variants: NA<br>Traits: NA | □ | ■ | □ | □ |
| Genomic range (chromosome, and, start and end positions) | Studies: NA<br>Associations: NA<br>E■ P■ Variants: /singleNucleotidePolymorphisms/search/findByChromBpLocationRange?chrom=22&bpStart=1&bpEnd=15500000<br>Traits: NA | □ | □ | ■ | □ |
| Gene name | Studies: NA<br>Associations: NA<br>E■ P■ Variants: /singleNucleotidePolymorphisms/search/findByGene?geneName=BRCA1<br>Traits: NA | □ | □ | ■ | □ |
| EFO trait | E■ P■ Studies: /studies/search/findByEfoTrait?efoTrait=lung%20adenocarcinoma<br>E■ P□ Associations: /associations/search/findByEfoTrait?efoTrait=lung%20adenocarcinoma<br>E■ P□ Variants: /singleNucleotidePolymorphisms/search/findByEfoTrait?efoTrait=lung%20adenocarcinoma<br>E■ P■ Traits: /efoTraits/search/findByEfoTrait?trait=lung%20adenocarcinoma | ■ | ■ | ■ | ■ |
| Authors' reported trait | E■ P■ Studies: /studies/search/findByDiseaseTrait?diseaseTrait=Breast%20cancer<br>Associations: NA<br>E■ P□ Variants: /singleNucleotidePolymorphisms/search/findByDiseaseTrait?diseaseTrait=Breast%20cancer<br>Traits: NA | ■ | □ | ■ | □ |

**Table S1** | GWAS Catalog REST API URL endpoints. A JSON response can be embedded (E■) or not (E□), and paginated (P■) or not (P□). Full squares indicate that GWAS Catalog entities are retrievable by the search criterion, whereas outlined squares indicate otherwise. URL endpoints are shortened, i.e., the common prefix `https://www.ebi.ac.uk/gwas/rest/api/` is omitted for clarity.
