## Additional File 2 for "gwasrapidd: an R package to query, download and wrangle GWAS Catalog data"

### GWAS Catalog access with gwasrapidd

#### Introduction

The **GWAS Catalog** is a service provided by the EMBL-EBI and NHGRI that offers a manually curated and freely available database of published genome-wide association studies (GWAS).

The GWAS Catalog data provided by the **RESTful API** is organized around four core entities:

- **studies**
- **associations**
- **variants**
- **traits**

#### Get GWAS Catalog Entities

**gwasrapidd** facilitates the access to the Catalog via the RESTful API, allowing you to programmatically retrieve data directly into R. Each of the four entities is mapped to an S4 object of a class of the same name.

| GWAS CATALOG | RETRIEVAL FUNCTIONS | S4 CLASSES |
|---|---|---|
| 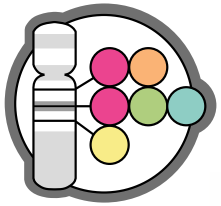 | `get_studies()` | S studies |
| | `get_associations()` | A associations |
| | `get_variants()` | V variants |
| | `get_traits()` | T traits |

| Search by | Example | S | A | V | T |
|---|---|---|---|---|---|
| study_id | "GCST000858" | ● | ● | ● | ● |
| association_id | "24300113" | ● | ● | ● | ● |
| variant_id | "rs12752552" | ● | ● | ● | ○ |
| efo_id | "EFO_0005543" | ● | ● | ● | ● |
| pubmed_id | "21626137" | ● | ● | ● | ● |
| user_requested | TRUE | ● | ○ | ○ | ○ |
| full_pvalue_set | FALSE | ● | ○ | ○ | ○ |
| efo_uri | "http://www.ebi.ac.uk/efo/EFO_0004761" | ● | ○ | ○ | ● |
| genomic_range | list(chromosome = "22", start = 1L, end = 15473564L) | ○ | ○ | ● | ○ |
| gene_name | "BRCA1" | ○ | ○ | ● | ○ |
| efo_trait | "lung adenocarcinoma" | ● | ● | ● | ● |
| reported_trait | "Breast cancer" | ● | ○ | ● | ○ |
| cytogenetic_band | "1p36.33" | ○ | ○ | ● | ○ |

#### S4 Representation of GWAS Catalog Entities

##### S4 class studies

The **studies** object consists of eight slots, each a table (tibble). Each study is an observation (row) in the `studies` table — main table. All tables have the column `study_id` as primary key.

For details about the studies S4 class: `class?studies`.

**studies**
- study_id
- reported_trait
- initial_sample_size
- replication_sample_size
- gxe
- gxg
- snp_count
- qualifier
- imputed
- pooled
- study_design_comment
- full_pvalue_set
- user_requested

**genotyping_techs**
- study_id
- genotyping technology

**platforms**
- study_id
- manufacturer

**ancestries**
- study_id
- ancestry_id
- type
- number_of_individuals

**ancestral_groups**
- study_id
- ancestry_id
- ancestral_group

**countries_of_recruitment**
- study_id
- ancestry_id
- country_name
- major_area
- region

**countries_of_origin**
- study_id
- ancestry_id
- country_name
- major_area
- region

**publications**
- study_id
- pubmed_id
- publication_date
- publication
- title
- author_fullname
- author_orcid

S

##### S4 class associations

The **associations** object consists of six slots, each a table (tibble). Each study is an observation (row) in the `associations` table — main table. All tables have the column `association_id` as primary key.

For details about the associations S4 class: `class?associations`.

**associations**
- association_id
- pvalue
- pvalue_description
- pvalue_mantissa
- pvalue_exponent
- multiple_snp_haplotype
- snp_interaction
- snp_type
- standard_error
- range
- or_per_copy_number
- beta_number
- beta_unit
- beta_direction
- beta_description
- last_mapping_date
- last_update_date

**loci**
- association_id
- locus_id
- haplotype_snp_count
- description

**risk_alleles**
- association_id
- locus_id
- variant_id
- risk_allele
- risk_frequency
- genome_wide
- limited_list

**genes**
- association_id
- locus_id
- gene_name

**ensembl_ids**
- association_id
- locus_id
- gene_name
- ensembl_id

**entrez_ids**
- association_id
- locus_id
- gene_name
- entrez_id

A

##### S4 class variants

The **variants** object consists of four slots, each a table (tibble). Each variant is an observation (row) in the `variants` table — main table. All tables have the column `variant_id` as primary key.

For details about the variants S4 class: `class?variants`.

**variants**
- variant_id
- merged
- functional_class
- chromosome_name
- chromosome_position
- chromosome_region
- last_update_date

**genomic_contexts**
- variant_id
- gene_name
- chromosome_name
- chromosome_position
- distance
- is_closest_gene
- is_intergenic
- is_upstream
- is_downstream
- source
- mapping_method

**ensembl_ids**
- variant_id
- gene_name
- ensembl_id

**entrez_ids**
- variant_id
- gene_name
- entrez_id

V

##### S4 class traits

The **traits** object consists of one slot only, a table (tibble) of GWAS Catalog EFO traits. Each EFO trait is an observation (row) in the traits table — main table.

For details about the traits S4 class: `class?traits`.

**traits**
- efo_id
- trait
- uri

T

#### Manipulate Cases

Get a **studies** object **s** of two GWAS studies:

```
s <- get_studies(study_id = c('a', 'b'))
```

Subset object **s** by either identifier or position using `` `[` ``:

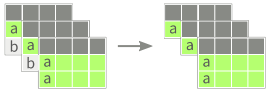

```
s['a'] # Subset by identifier
```

```
s[1] # Subset by position
```

Combine two studies' objects:

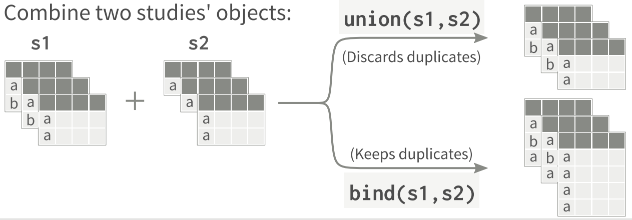


CC BY SA Ramiro Magno • Learn more at https://rmagno.eu/gwasrapidd • gwasrapidd version 0.0.0.9014 • Updated: 2019-04-28
